# scSTATseq: Diminishing Technical Dropout Enables Core Transcriptome Recovery and Comprehensive Single-cell Trajectory Mapping

**DOI:** 10.1101/2020.04.15.042408

**Authors:** Zihan Zheng, Xiangyu Tang, Xin Qiu, Hao Xu, Haiyang Wu, Haili Yu, Xingzhao Wen, Zhou Peng, Fa Xu, Yiwen Zhou, Qingshan Ni, Jianzhi Zhou, Liyun Zou, Gang Chen, Ying Wan

**Author notes:** These authors contributed equally to this work (Zihan Zheng, Xiangyu Tang, Xin Qiu).

## Abstract

The advent of single-cell RNA sequencing has provided illuminating information on complex systems. However, large numbers of genes tend to be scarcely detected in common scRNAseq approaches due to technical dropout. Although bioinformatics approaches have been developed to approximate true expression profiles, assess the dropout events on single-cell transcriptomes is still consequently challenging. In this report, we present a new plate-based method for scRNAseq that relies on Tn5 transposase to tagment cDNA following second strand synthesis. By utilizing pre-amplification tagmentation step, scSTATseq libraries are insulated against technical dropout, allowing for detailed analysis of gene-gene co-expression relationships and mapping of pathway trajectories. The entire scSTATseq library construction workflow can be completed in 7 hours, and recover transcriptome information on up to 8,000 protein-coding genes. Investigation of osteoclast differentiation using this workflow allowed us to identify novel markers of interest such as Rab15. Overall, scSTATseq is an efficient and economical method for scRNAseq that compares favorably with existing workflows.

## Main

Single cell RNAseq has become a highly useful tool for interrogating biological systems, and permitting finer understanding of the transitory cellular states following stimulation in both homogenous and heterogeneous populations. A number of different methods have been developed to generate scRNAseq libraries, each with distinctive advantages^1^. Droplet-based approaches such as inDrop^2^ and Microwell-seq^3^ have emphasized capture of large amount of cells at the expense of individual cell detail. In contrast, commonly used plate-based approaches, such as SMARTseq2^4^ and CEL-seq^5^, can generate highly detailed profiles of small numbers of cells, including more information on splicing events and non-coding sequences.

However, despite their divergent protocols, all of these methods share a requirement for an initial amplification of RNA content, either through whole transcriptome amplification (WTA) or in vitro transcription (IVT), under the presumption that the initial cDNA quantity of any given cell is too minute to work with. However, biases in the transcriptome pool may result from the initial amplification, as some cDNA evade amplification/mRNA capture^6^. These biases contribute to technical dropout, wherein uneven and pseudo-random detection of medium- and low-expressed genes significantly occludes scRNAseq results and has been recognized as a key concern for single-cell experiments^7,8,9^. While a number of computational approaches have been designed to help overcome dropout through imputation, such methods have difficulty in recovering true gene-gene co-expression relationships and consequent co-expression networks^10,11^.

In order to streamline scRNAseq library construction and lower technical dropout, we hypothesized that early fragmentation of the cDNA prior to large-scale amplification would help avoid transcript biases, akin to the RNA fragmentation approach used in some bulk protocols. As such, in the scSTATseq workflow presented here (Fig 1A), sorted cells are rapidly lysed, and subsequently reverse transcribed into cDNA with the use of conventional oligo-dT, together with not-so-random hexamers (NSR) designed to help capture 5’ information^12^. A rapid second-strand synthesis step is used to generate paired cDNA strands, which are then immediately tagmented via homebrew Tn5, such that cDNA amplification only occurs after fragmentation is completed. Samples are only pooled together following the completion of sequencing index ligation, and sample transfer steps were minimized, in order to prevent potential cross-contamination.

**Figure 1.**
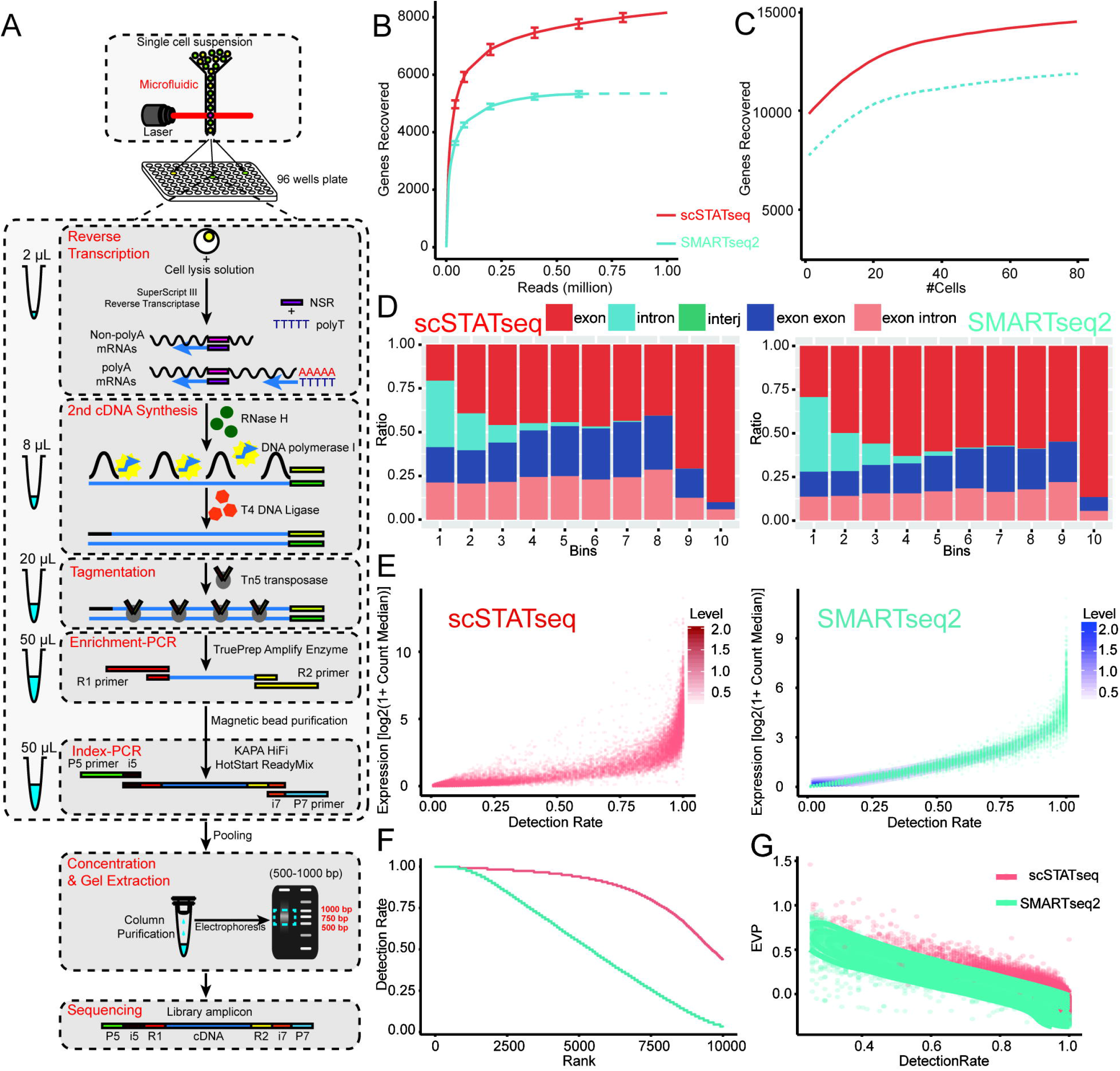
scSTATseq workflow enhances transcriptome recovery. **A)** Overview of the scSTATseq workflow (a detailed protocol is included in the methods section). **B)** Saturation curves of the number of protein-coding genes found from 80 scSTATseq libraries or a publicly available dataset of SMARTseq2 libraries (80 control RAW cells) as a function of sequencing depth. Although scSTATseq libraries were sequenced to a greater depth than the SMARTseq2 sets, the SMARTseq2 set appeared to have reached saturation at around 0.5 million reads/cell, while the scSTATseq2 libraries detected an average of over 2,000 more genes at the same depth. **C)** Saturation curves for the amount of genes detected as a function of increasing numbers of cells sequenced. Since both datasets were generated using a stable cell line, the genes detected in any given cell are highly similar, with sequencing of 40 cells being sufficient to reach saturation for commonly expressed genes. **D)** Examination of the genomic mapping of the raw scSTATseq and SMARTseq2 libraries binned into quantiles based on expression level (1 lowest, 10 highest). While a steady increase in the proportion of intronic reads can be found in the lower bins of both scSTATseq and SMARTseq2 libraries, indicative of potential genomic DNA contamination among rarer transcripts, scSTATseq maintains much higher rates of exon-exon spanning reads throughout. **E)** Scatterplot comparison of detection rate of scSTATseq and SMARTseq2 libraries, downsampled to 0.5 million reads/cell as a function of gene expression level. Although both scSTATseq and SMARTseq2 libraries include a substantial number of low-expressed genes that are found in only a small portion of cells, the scSTATseq libraries had a much lower proportion of cells with intermediate expression and detection (∼50% non-zero detection). The preservation of expression information for these genes reflects a lowered rate of technical dropout. **F)** Summary of dropout rates in the downsampled libraries (as in D) ranked from highest (feature found in every cell) to lowest (feature uncommonly found in cells). While the technical dropout rate in the SMARTseq2 data exhibited a sharp linear decrease, with only a small pool of 796 features being found in every cell and 3,299 features found in >75% of cells, scSTATseq recovered 2,781 features across all cells. 577 of the 796 features found in each of the SMARTseq2 cells were also found in each scSTATseq cell, while 180 of the remainder were found in >90% of the scSTATseq cells. At the same time, 839 other common features in scSTATseq cells could be found in >90% of the SMARTseq2 cells. This high overall similarity in features suggests that the difference in detection rate is the product of technical dropout, and not likely the result of sharp differences between the RAW cell starting material used. **G)** Scatterplot of the extra-Poisson variation for each given gene in the SMARTseq2 and scSTATseq libraries as a function of its detection rate. Genes detected in less than 25% of the cells in either library were excluded. SMARTseq2 has higher levels of EPV overall as a result of a long tail of genes with low expression, despite having low levels of variation in genes with high detection. See also Fig S3B-C.

The entire workflow can be completed by hand in 7 hours, with favorable per-cell cost compared to other common plate-based workflows (Table S1). From our initial applications of scSTATseq to the mouse RAW264.7 macrophage cell line, we could recover a median of just under 9,000 protein coding genes per cell at a sequencing depth approaching 10million reads/cell across 160 cells (Fig S1A) and with a majority of genes displaying medium expression levels (Fig S1B). Similar numbers of genes were found in each batch of cells, with high genomic mapping ratios and relatively low percentages of mitochondrial and rRNA reads (Fig S1C). These cells could be stratified based on cell cycle stage, with both S phase and G2-M phase cells identifiable in each batch (Fig S2A-D). Information on other types of non-coding sequences, such as microRNA housing genes and lncRNAs, could also be recovered (Fig S2F-H). To assess the robustness of scSTATseq, we then compared the RAW cell profiles obtained using our approach with a publicly available dataset generated according to the SMARTseq2 protocol^13^. Analysis of saturation curves of both libraries showed that scSTATseq could recover transcriptome information on 3,000 more protein coding genes than SMARTseq2 on average, an improvement independent of sequencing depth or cell count (Fig 1B-C). These uniquely detected genes tended to have lower median expression than their SMARTseq2-shared counterparts (Fig S3A). However, inspection of the genomic mapping revealed that these low-expressed genes in the scSTATseq data still maintained a high rate of exon-intron and exon-exon spanning reads, indicative of successful immature and mature mRNA detection (Fig 1G).

In order to assess the significance of the higher numbers of genes recovered, we then inspected the detection rate and expression distributions of the two datasets. We observed that genes in the scSTATseq data displayed a bimodal detection pattern that was influenced by expression level (a prominent peak of genes broadly detected in >75% of cells and with high average expression, with a sparser number of genes with detection rates between 25% and 75%), while the peaks in the SMARTseq2 detection distribution were much less distinct (such that most of the medium-expressed genes had dropout rates > 25%, with a dense linear correlation between expression level and detection rate) (Fig 1D). While the observation that low-expressing genes are more prone to dropout is of itself unsurprising, the even linearity of the correlation in the SMARTseq2 data suggests that a significant portion of the dropout is the product of technical variation (which would be biased in favor of keeping more highly expressed genes), and not transcriptional burst dynamics (which should be more random). As such, the distributional difference suggested to us that scSTATseq libraries were able to more efficiently preserve higher detection rates for medium and low expressing genes by diminishing some dropout caused by technical variation. Consistent with this expectation, we observed that genes with higher dropout displayed higher degrees of extra-Poisson variation (EPV), and the scSTATseq data consequently had higher overall precision as a result of having lowered dropout (Fig 1F, S3B-C). Taken together, these results demonstrate that scSTATseq libraries are significantly insulated against zero-inflated dropout.

To further characterize the consequence of the reduction in dropout on downstream analyses, we next performed co-expression analysis using both datasets to identify the genes with the highest conservation. Although this type of analysis is commonly seen for bulk-sequencing studies, high dropout rates and small dynamic ranges have made their single-cell implementation difficult. Consistent with this understanding, the numbers of co-expressed pairs was quite low in both types of single-cell libraries considered (Fig 2A). However, the scSTATseq data showed significant numbers of gene pairs that had higher correlations (*R >* 0.4), with the background correlation in both sets being below 0.2 (Fig S3D). In order to further clarify the contribution of technical dropout to this divergence in co-expression, we then evaluated the dropout rate of the genes pairs which were coexpressed in one library but not the other. Interestingly, 78% of the scSTATseq correlates were detected at similar levels in both libraries (detection difference less than 20%), while only 22% of the pairs showed heavy dropout in the SMARTseq2 data (Fig 2B). Examination of the overall gene expression distribution of both libraries demonstrated that the scSTATseq data was more platykurtic and was not noticeably skewed, while the SMARTseq2 data displayed a rightward skew (Fig S3E). At the same time, the scSTATseq data showed higher dispersion among genes at the lower and higher ends for expression, indicating insulation against threshold effects (Fig S3F). Collectively, these results demonstrate that scSTATseq has both a larger effective dynamic range for expression detection and lower rates of technical dropout, rendering it more amenable for co-expression analysis.

**Figure 2.**
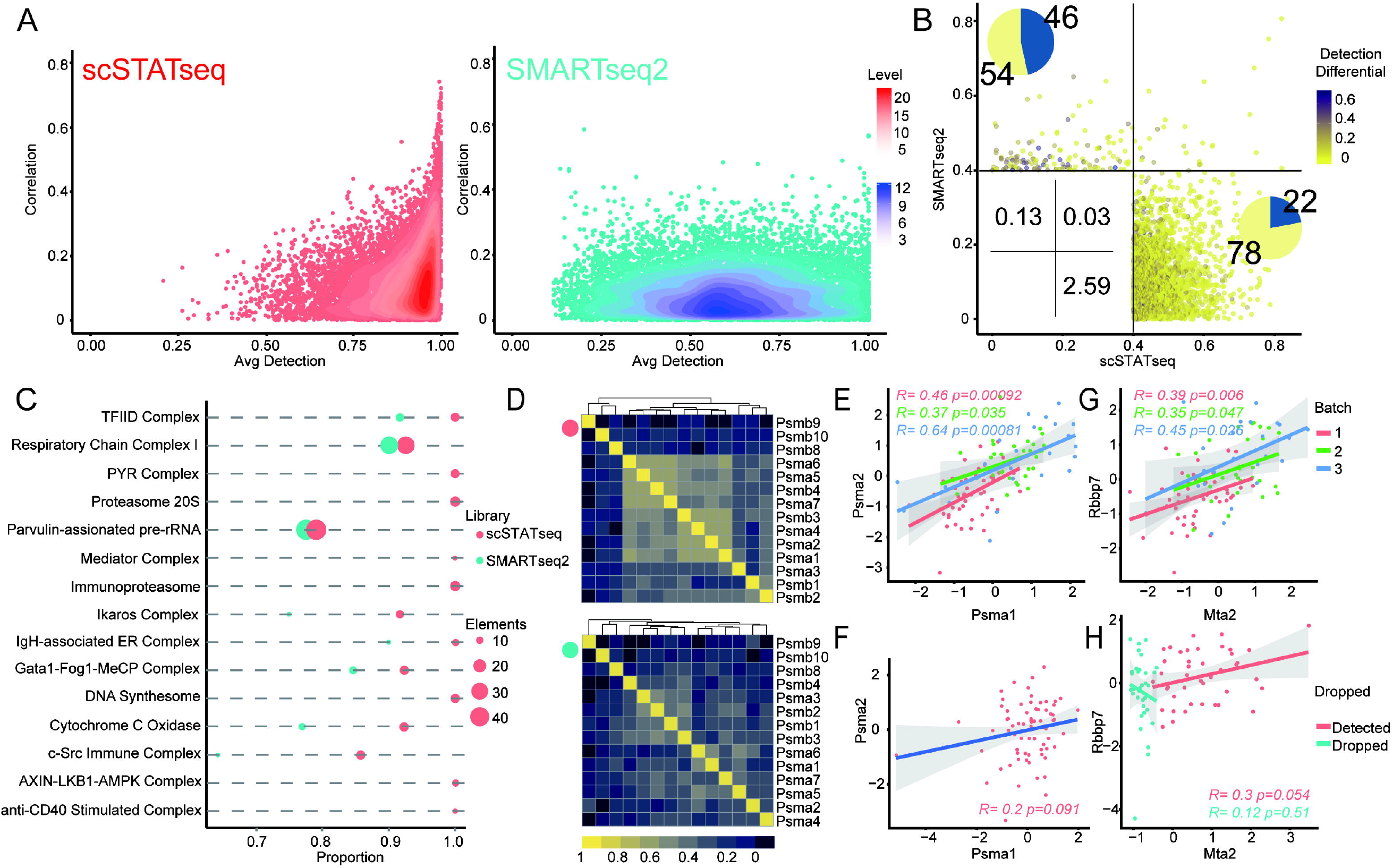
Identification of a stable transcriptome through co-expression analysis. **A)** Pairwise absolute Pearson correlation of 10,000 randomly selected pairs of genes plotted against the averaged detection rates for each gene pair. Genes detected in less than 10% of either library type were excluded to permit a more direct examination of the most conserved genes. A total of 269 pairs in the scSTATseq libraries had significant correlations (*R* > 0.4) and the minimum average detection rate for these pairs was 75%, while 23 pairs could be found in the SMARTseq2 data with minimum average detection rate of 20%. **B)** Expanded pairwise correlation analysis of 100,000 gene pairs in both SMARTseq2 and scSTATseq libraries. Absolute Pearson correlation values for a given gene pair in scSTATseq (x-axis) and SMARTseq2 (y-axis) libraries are plotted together to characterize the differences in correlation. Of note, the distribution of the scSTATseq-unique correlates (2.59% of the pairs seen in quadrant IV) are skewed towards a SMARTseq2-correlation of 0 and not towards a SMARTseq2-correlation close to 0.4 (overall skew of 0.91 and kurtosis of 0.28), demonstrating that the difference was not due a multitude of pairs being on the edge of the threshold in the SMARTseq2 data. Each pair is colored based on absolute detection differences between the two libraries. 22% of the correlates found in the scSTATseq libraries showed significantly lower detection rates in the SMARTseq2 data, and their lack of detection may be directly attributable to technical dropout in SMARTseq2 (see also SF3D). **C)** Dotplot of the number of unique elements of murine protein complexes (from the CORUM database) with significant expression correlation in either scSTATseq or SMARTseq2 libraries. The smallest complexes (< 5 elements) were excluded from the plot to prevent overrepresentation, but a full table summarizing the results is included as Table S). **D)** Correlation heatmaps for the elements in the immunoproteasome complex in both scSTATseq and SMARTseq2 data. While 8 elements are strongly correlated in the scSTATseq data, only scattered pairwise correlations can be seen in the SMARTseq2 data. **E)** Closer examination of the correlation between the immunoproteasome components *Psma1* and *Psma2* in the scSTATseq data demonstrates that the correlation between these two proteins is statistically significant across three independent batches of cells. **F)** Similar examination of *Psma1* and *Psma2* in the SMARTseq2 data fails to recover a significant correlation in expression. Notably, neither gene had heavy dropout in the SMARTseq2 data, suggesting that the lack of correlation was independent of dropout and more likely a product of variance in signal range. **G)** The Gata1-complex members *Mta2* and *Rbbp7* are significantly co-expressed across all three batches of the scSTATseq data. **H)** No correlation can be seen between these two genes in the SMARTseq2 data, as a result of heavy dropout in the detection of *Mta2* (missing values in 46% of the cells). This is a representative demonstration of the loss of co-expression as a direct result of technical dropout.

In order to evaluate the functional importance of the genes found to be co-expressed in the scSTATseq data, we next leveraged existing databases for protein-protein interaction (CORUM) and functional pathways (KEGG) for further analyses. The scSTATseq data was able to detect both larger numbers of genes in the CORUM database (Fig S3), and also detect high levels of intercorrelation in a larger number of complexes (Fig 2C, Table S2). These complexes include ones known to be critical for macrophage function, such as the immunoproteasome complex responsible for enabling antigen presentation, as well as ones less well characterized, such as one centered on the transcription factor Gata1 (Fig 2D). Direct inspection of the highly correlated Psma1-Psma2 immunoproteasome gene pair demonstrated that a positive correlation in expression could be seen across all three replicates of the scSTATseq data, but was lost in the SMARTseq2 data due to its narrow detection of Psma1 expression, even though it was positively detected across all cells in both libraries (Fig 2E). At the same time, no correlation in the expression of Mta2-Rbbp7 could be found in the SMARTseq2 data due to heavy dropout in the expression of Mta2 (Fig 2F). Similar results were obtained from analysis using KEGG pathways (Fig S4, Table S3). These results thus informed us that the scSTATseq libraries preserved significantly greater numbers of correlated gene pairs with known biological significance.

Having confirmed the robustness of co-expression pairing on the scSTATseq data, we then sought to identify the core transcriptome of RAW cells on a single-cell level by screening for the genes that had high detection, high expression, low variability, and which were highly inter-correlated with other genes. Comprehensive screening and network generation based on these parameters yielded a pool of 3,675 genes, representing just over 10% of all features detected (Fig S5A, Table S4). Each of these genes was significantly correlated with over 180 other genes on average (Fig S5B), and community clustering identified 5 primary clusters of genes (Fig S5C-D). These clusters were enriched for genes with distinct annotated biological functions (Fig S5E, Table S5). Interestingly, one of these clusters displayed prominent enrichment of immune-related pathways (Fig S6A). Direct inspection of the cluster demonstrated that a number of genes with well-characterized immune functions (such as *Ptprc*, *Lyz2*, and *Tlr2*) could be found within, while other genes with less established roles were found to be significantly linked (Fig S6E). The identification of this core transcriptome may be useful as a background reference list of essential macrophage genes and for screening novel molecular mechanisms involved in macrophage function.

We next sought to exploit these advantages of scSTATseq to investigate a dynamic process that could be modeled using RAW cells, namely osteoclast differentiation. Although osteoclasts are a primary pathogenic cell type contributing to osteoporosis, the molecular mechanisms underlying their differentiation remain incompletely understood, and may be of interest for therapeutic development^14,15,16^. Sequencing of RANKL-stimulated RAW cells at 4 time points revealed a similar set of genes recovered and low dropout rates across all samples (Fig S7), with clear time-dependent changes in the expression of a number of factors previously associated with osteoclast differentiation (Fig 3A). In order to map out the trajectory of osteoclast differentiation, we first constructed a baseline pseudotime trajectory (Fig 3B), wherein the 48 and 72 hr cells had the highest pseudotime progression (Fig 3B) following progression through some minor intermediate states (Fig S8). We then computed trajectories and assigned pseudotime values for progression along 2,263 curated pathways from Reactome. Via correlation analysis of pathway-specific pseudotime values against the baseline pseudotime values, we could then infer the association of the pathway with the overall progression trajectory. From this analysis, we recovered 26 pathways with high absolute correlation, including vesicular transport and interleukin signaling (Fig 3C, Table S5).

**Figure 3.**
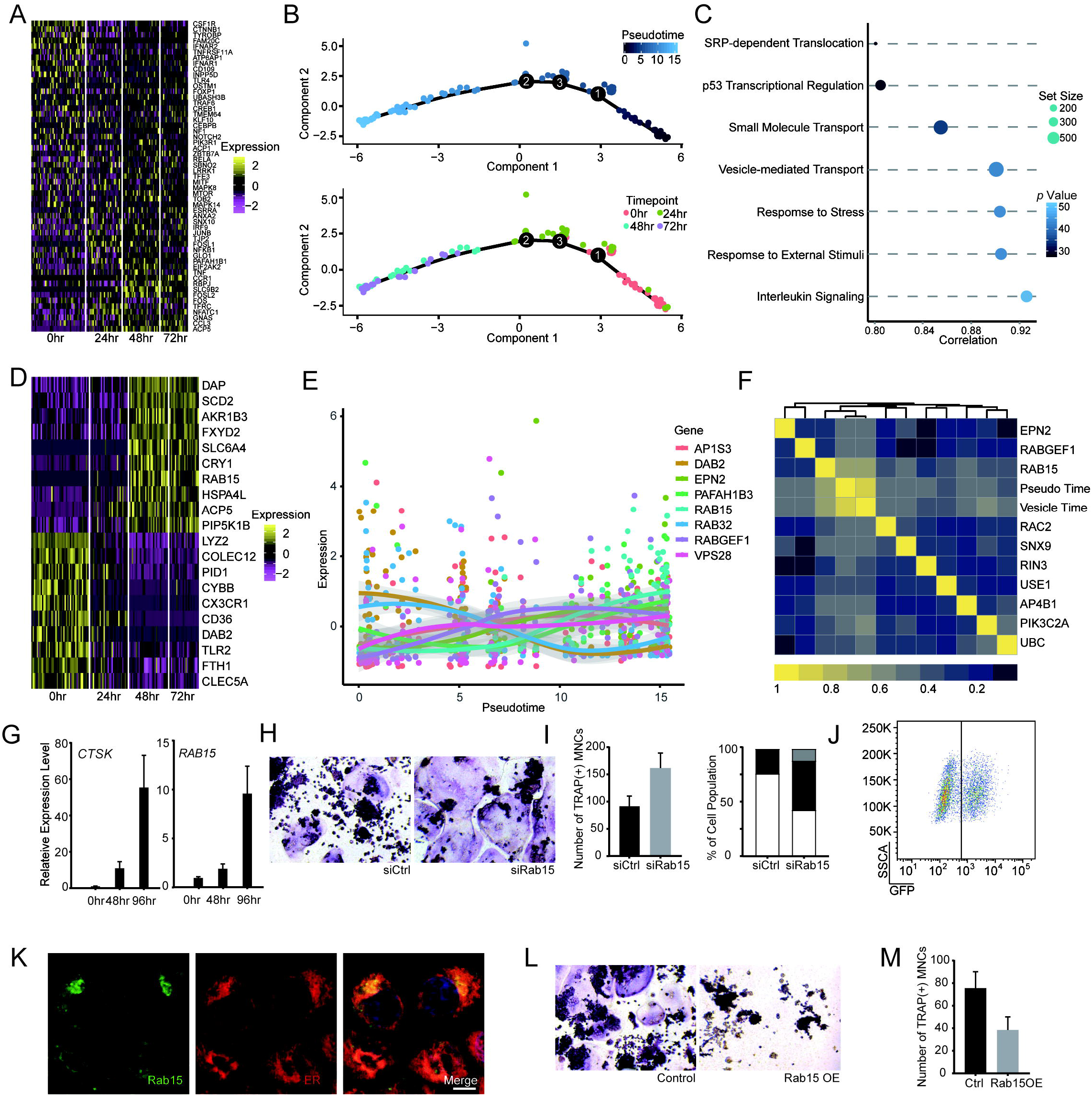
scSTATseq analysis of osteoclast differentiation. **A)** Heatmap of the 55 genes from the KEGG curated osteoclast differentiation pathway with significant changes in expression between 0hr and 72hr samples across the 122 cells sequenced. Cells were collected simultaneously to limit batch effects, with cells of two timepoints paired onto the same plate to further rule out the possibility of high levels of similarity being driven by cross-contamination. **B)** Pseudotime trajectory of osteoclast differentiation computed using DDRTree dimension reduction within Monocle using 823 highly-variable genes. While median pseudotime values for the 48 and 72 hour cells were indistinguishable, the 24 hour cells showed significant progression from baseline controls. **C)** Dot plot of the pathway trajectories most highly correlated with the overall osteoclast differentiation trajectory. Correlations are given as absolute Pearson’s R since the actual directionality of pseudotime assignment is not necessarily significant. **D)** Heatmap of the top10 most positively and negatively correlated genes with the pseudotime trajectory defined in (B). Notably, while 4 of these factors were contained in the variable gene list used to construct the trajectory, the other factors were not, demonstrating that the trajectory analysis could recover information on co-expressed genes. **E)** Expression level of the vesicle-transport genes displaying switch-like behavior over the primary pseudotime trajectory. While a number of these factors display their switch early on in the pseudotime trajectory, *Rab15* and *Pafah1b3* switch at the boundary of the change between 24 to 48/72hr samples. **F)** Heatmap of the correlations between proteins involved in vesicular trafficking and the overall pseudotime trajectory and vesicular trafficking trajectory. Notably, while a number of genes display a statistically significant level of correlation, the relationship with *Rab15* stands out. **G)** qPCR of the expression of Ctsk and Rab15 in RAW cells stimulated for 0, 48, and 96 hours confirms that Rab15 mRNA expression appreciably increases at 48hr and peaks at 96hr. **H)** TRAP stain of 96hr-stimulated Raw cells identifies larger numbers of fused mature osteoclasts following Rab15 depletion through siRNA transfection. **I)** Depletion of Rab15 causes a larger proportion of large osteoclasts to form, leading to an emergence of a population of giant osteoclasts with >50 nuclei. **J)** Transfection of a Rab15-GFP fusion plasmid in RAW cells allows for selection of an overexpressing population. **K)** A portion of the overexpressed Rab15 appears to localize in the endoplasmic reticulum. **L-M)** TRAP stain of 96hr-stimulated RAW cells demonstrates that Rab15 overexpression greatly suppresses the differentiation of mature osteoclasts.

Since the general importance of vesicular transport to osteoclast differentiation has been previously demonstrated in the context of enzymes such as CTSK and ACP5^17^, we were unsurprised to see it correlate with the overall pseudotime progression. However, the contribution of individual proteins responsible for transporting these vesicles remain incompletely understood. As such, we then searched for gene-switching events within the trafficking-associated pathways to identify the key molecules involved (Fig S9, Table S6). From this analysis, we found that the ras-related protein *Rab15* was switched on in the 48-72hr samples (Fig 3D), and traditional correlation analysis confirmed that *Rab15* expression was also significantly correlated with overall trajectory progression (Fig 3E). While Rab15 was not previously identified in screens of progenitor cells^18^, careful inspection of the reads mapping to Rab15 confirmed a strong positive signal on the last exon (Fig S10E), and its expression could also be validated via qPCR (Fig 3G) and immunofluorescence (Fig 3H). As such, we then used overexpression and knockdown experiments to assess its influence. Surprisingly, overexpression of Rab15 drove a significant decrease in mature osteoclast formation (Fig 3I). At the same time, Rab15 depletion led to an increase in osteoclast differentiation, with a sharp increase in large, TRAP+ fused osteoclasts (Fig 3J).

Given that Rab15 has been previously demonstrated to influence trafficking from early endosomes, this behavior suggests that osteoclasts may also negatively regulate their own function through differential processing of newly endocytosed receptors^19^. Although the exact identity of the cargo remains unclear, it nonetheless validates the importance of the vesicular transport pathway identified through our analyses. Furthermore, while many positive regulators of osteoclast differentiation have been previously identified, clear negative regulators are much less commonly observed^20^. Identification of these negative regulators as a result of the increased clarity offered by scRNAseq may thus expand our understanding of the entire process and uncover new therapeutic targets.

By streamlining a scRNAseq workflow and performing cDNA fragmentation prior to amplification, we were able to observe a clear improvement in both protein-coding gene count and detection rate. While concerns regarding the minute amount of starting material have led most workflows to develop early amplification steps, we found that second-strand synthesis alone could provide a sufficient base for later enrichment, especially since we maintained each reaction in individual wells and only added new reagents. Indeed, our initial concerns about potentially high amounts of contaminating gDNA being amplified by this process proved to be unfounded, as the majority of reads consistently mapped to exon regions even in genes with relatively low expression. Instead, rapid tagmentation of the transcribed cDNA seemed to expand the dynamic range for expression detection. The consequent improvement in data resolution permits enhanced interrogation of true gene co-expression relationships in single-cell data and more detailed investigation of transcriptional regulation patterns. Use of the scSTATseq workflow also greatly improves the resolution of trajectory mapping analysis and may help illuminate studies of complex cellular differentiation mechanisms.

## Acknowledgements

The authors would like to thank the other members of the lab for their helpful suggestions during the development of this method. This work was supported in part by grants from the National Key Research and Development Program of China (2017YFA0700404) and National Natural Science Foundation of China (91642119) Y.W., Basic Science and Frontier Technology Research Project of Chongqing (cstc2017jcyjAX0198) to C.G., and National Natural Science Foundation of China (81971546) to L.Z.

---

**Figure S1-**scSTATseq quality control and clustering

A) Three batches of RAW cells were prepared separately and merged together for the analyses in Fig 1 had similar median protein coding gene and pseudogene counts. Outlier cells with over 12,000 protein coding genes were excluded as potential doublets/dead cell artifacts. B) Histogram of gene count ordered by expression level (log2 transcripts per million reads) shows a peak of genes between 3 and 8 TPM comprise the majority of the features detected in the libraries (complementing Fig 1D). C) Transcript mapping ratio to mouse genome across all three batches showed low rates of primer dimers or other contaminating sequences. Cells with mapping rates below 75% were excluded due to possible contamination. While mitochondrial RNA ratio is often used as a quality control for droplet-based datasets, we elected not to apply it as a strict filter for scSTATseq libraries due to the differences in method. Instead, an upper threshold of 20% mitochondrial RNA was set in the interest of excluding cells that would otherwise have too little transcript information. Similarly, we applied a rRNA ratio filter of 30% to exclude cells dominated by rRNA reads.

**Figure S2-**Cell cycle clustering of RAW cells

A) UMAP reduction of three batches of RAW cells based on a list of known cell cycle genes (from Tirosh et al. 2016). Cells from all three batches are generally admixed, suggesting an overall similarity in expression profile for these genes across the three batches. B) UMAP visualization of the expression of several prominent cell cycle markers showing differential expression across the UMAP space (*Mki67* expression is higher in the upper left cells, while *Pcna* expression is higher in the lower right). C) Overall cell cycle scoring for each cell demonstrates that the cells in the lower right are predominantly in S phase, while those in the upper left are in G2-M phases. D) Frequency of the cells belonging to each cell cycle stage across the three batches. Cells with score > 10 for either G2-M or S phase were scored as belonging to those phases, while cells with >10 score for both scores were deemed as intermediates. Cells with <10 score for both were labeled as quiescent. While there is substantial variability in cell cycle across the three batches, cells in each class can be found in every batch. E) Expression of four functional molecules commonly found on macrophages (and the lack thereof) suggests that the RAW cells were resting and had not undergone activation (Cd69-across all cells, and rare expression of Pd-l1 and Cd86). F) Since the libraries contained large amounts of pseudogenes, we further investigated if scSTATseq may provide some information on long non-coding sequences. UMAP visualization of two commonly expressed lncRNAs (*Malat1* and *Neat1*) showed that both had relatively high expression, with some enrichment for these two lncRNAs in the cluster of cells with higher S phase scores. G) At the same time, while microRNAs are too short to be successfully sequenced by scSTATseq, we observed that the expression of several microRNA housing genes could be recovered from the libraries, and may provide some insight into mature microRNA expression. H) We further examined the well characterized lncRNAs in the X-inactivation system as an additional check. Neither the primary chromosome-coating Xist nor its antisense transcript Tsix were found to be significantly expressed. On the other hand, some expression of known escape genes Jpx and Ftx could be found, albeit at relatively low levels. Since RAW 264.7 cells are derived from a male BALB/c mouse, this pattern is consistent with our expectations.

**Figure S3-**Broad correlation and distribution comparison

A) Histogram visualization of the median expression levels of the 577 features in scSTATseq data that are either shared between the two datasets (blue) or those uniquely found at high rates in scSTATseq libraries (salmon) shows that the uniquely detected features are of significantly lower median expression level. B) Comparison of the extra-Poisson variation in the scSTATseq and SMARTseq2 libraries for a given gene. Genes found in less than 25% of the cells in either library type were excluded. Most of the high-EPV genes in the SMARTseq2 data fell into a narrow EPV distribution in the scSTATseq data, albeit with a slight rightwards skew. C) Summary of the EPV distribution of the scSTATseq and SMARTseq2 libraries. The scSTATseq EPVs fell into a narrow, leptokurtic distribution while the SMARTseq2 distribution is much broader, suggesting that the technical precision of scSTATseq is significantly higher for a large number of genes. D) Summary of the summed expression of each gene in the two libraries, post-scaling. The expression in the SMARTseq2 is noticeably flatter and slightly skewed to the left, while scSTATseq pattern is more balanced and features a slightly longer linear range. E) Scatterplot of standard deviation in relation to gene expression. While the SMARTseq2 data displays prominent decreases in SD at both lower and upper expression thresholds (suggesting strong influence of upper and lower bounds of detection), this decrease is more subtle in the scSTATseq data due to its wider dynamic range. F) The number of gene pairs identified in either scSTATseq or SMARTseq2 data at different correlation thresholds (100* Pearson’s R). A thousand lists of 100 randomly selected genes chosen from the pool of common genes between the two methods was used for calculation, allowing for a maximum of 4950 unique pairs. Notably, at a correlation level of R > 0.4, a median of 124 genes were found, representing 2.7% of all possible pairs. This proportion of correlated pairs is essentially identical to our observation in Fig 2B, consistent with our expectations from a random selection. We thus established a R > 0.4 as the threshold and representation of 2.7% for the lower baseline to determine whether a given gene set displayed true overrepresentation of correlated pairs. G) Comparison of the number of elements from a given CORUM complex that can be found in either library type. H) Average detection rates for all of the detected elements in a CORUM complex in either library type. Detection values tend to be much higher in the scSTATseq data, with a large patch of complexes showing >20% higher mean detection rate. I) scSTATseq data allows for observation of correlations in some cases where no correlates are found in SMARTseq2. In the PYR complex example shown, the SMARTseq2 data has no pairwise correlations >0.4, and only scattered correlations >0.2. However, the scSTATseq data shows 12 pairs of correlates >0.4.

**Figure S4-**Correlated expression of KEGG pathways in scSTATseq data

A) Comparison of the number of elements from a given KEGG pathway that can be found in either library type. Overall, the scSTATseq libraries showed much higher proportional representation as a result of the higher numbers of gene detected. B) Median detection rates of the elements recovered from a given pathway in either library type. Similar to the observation in Fig S3H, scSTATseq featured higher pathway element detection rates. C) While the SMARTseq2 data featured more ribosome elements that displayed pairwise correlation, the scSTATseq data showed higher numbers of correlates in pathways critical for macrophage function, such as proteasome, leukocyte transendothelial migration, and chemokine signaling. D) Heatmaps showing correlations in both library types of elements in the antigen processing and presentation pathway. Since one of the key functions of macrophages is to act as professional antigen-presenting cells, it is unsurprising that many of its elements may have highly similar expression profiles. However, these relationships are not clearly captured in the SMARTseq2 data. E) Heatmap showing correlations in both library types of elements in the larger lysosome pathway. Similar to the results in the smaller sets, a clear cluster of highly correlated genes can be found in the scSTATseq data, including an assortment of cathepsins (including *Ctsa, Ctsb, Ctsd,* and *Ctsz*, among others), as well as the proton pump *Atp6v0b* and key marker *Lamp1*.

**Figure S5-**Core transcriptome of RAW cells

A) Network visualization of the gene-gene correlations in RAW cells. Pair-wise correlations of R > 0.5 are retained as edges, with higher correlations matching to darker colors. Size of gene name label corresponds to the degree of the given node (larger nodes are ones with more correlation relationships). B) Distribution of the number of connections for each node. Median degree for all nodes is 181, demonstrating that the overall network is rather extensively interconnected. C) Clustering of the network at resolution of 1 identifies 27 clusters, of which 5 (12, 23, 24, 25, and 26) include a larger number of genes. D) Edges recolored by their source to demonstrate the spatial locations for each of the five larger clusters of genes identified. E) Gene set overrepresentation analysis to identify the prominent Reactome pathways that genes from each of the clusters belong to. While cluster12 shows prominent enrichment for cell-cycle genes, (foldenrich calculated as number of genes found over expectated based on gene set size), cluster23 appears to include a number of genes involved in immune cell function, such as TLR recycling and free radical production.

**Figure S6**-Inspection of immune-related clsuter23

A) Close-up visualization of the genes in cluster23 shows clear inclusion of a number of factors with known participation in immune function, such as the integrins *Itgb2* and *Itgav*, as well assorted surface molecules such as *Tfrc*, *Ptprc*, and *Sirpa*. Notably, a number of orange and pink-colored genes can be discerned in the close-up. While it has been demonstrated that genes with higher expression levels tend to have higher degrees of expression correlation, the inclusion of these genes with lower expression levels suggests the network constructed is not entirely limited by expression. B) Similar to the distribution in the overall network, gene nodes in cluster23 are also highly interconnected. C) Violin plot of the number of connected nodes to each gene, stratified based on expression level, offering a statistical demonstration of the observation described in (A). D) Demonstration of the high degree of connectivity for nodes in the cluster23 network. Using two randomly selected points as the starting point, we could observe that the majority of the other nodes were within a 2-neighbor distance of the starting point. E) Clarification of the core transcriptome can help identify other genes that may be involved in relevant biological processes using guilt-by-association. Genes which are correlated across multiple batches as shown here may be promising candidates for further biological validation.

**Figure S7-**QC of osteoclast single cell sequencing

A) A consistent number of genes were found in the cells across all four timepoints. Rare doublets with more than 15,000 genes and lower-quality cells with fewer than 5,000 genes were excluded from further analysis. B) Evaluation of dropout rates across all four timepoints. As some factors may be restricted to a particular timepoint, we did not rely on a shared pool of genes, but instead considered each timepoint individually and ranked all genes found in a given timepoint by their detection rate. C) Intersection analysis of the genes found across the four timepoints demonstrates that a larger majority of genes (5,717) are found in all four timepoints, although a number of unique factors (606) were found in the 48/72hr samples, suggesting that these may encompass some osteoclast-specific inducible factors. D) Comparison of the number of elements found in CORUM complexes between the 72hr and 0hr samples found a small increase in detection of two complexes. E) UMAP reduction of the osteoclast data based on all elements of CORUM complexes shows that 48 and 72hr samples are largely intermixed, suggesting that protein complexes have their expression levels set by 48hrs following stimulation. F) Some of the CORUM complexes with significant numbers of correlated elements in the 48/72hr cells. G) Cell-cycle scoring of the osteoclasts (similar to Fig S2C). While a higher proportion of the 0/24hr cells are in S phase, suggesting a higher rate of replication, a number of the 48/72hr cells also appear to still be actively transitioning through the cell cycle. H) Violin plot of the scores in (G). The presence of these potentially replicating cells following 72hr of stimulation suggests that osteoclasts may not necessarily become quiescent during their differentiation.

**Figure S8**-Branching trajectory of osteoclast differentiation

A) Pseudotime trajectories generated using the list of KEGG curated osteoclast genes displayed in part in Fig 3E. A similar pattern can be seen where the 0hr samples mostly represent an earlier progression state, while the 25hr, 48hr, and 72hr samples all have some cells with highly advanced progression. B-C) Minimum spanning tree of the overall pseudotime trajectory computed in Fig 3D. While a total of 7 states can be seen within the tree, some states are minor nodes with relatively few cells from the 24hr sample. The greatest difference from the initial state in this context is state 4, comprised of the end-differentiated osteoclasts. D) Heatmap of the top markers for each of the individual states identified. Notably, the key osteoclast functional molecule *Ctsk* is prominently enriched in state 4, but is not found in the earlier states. E) UMAP dimension reduction of the osteoclast gene set (generated using the same list of highly variable genes that were considered for the original trajectory inference) shows clear separations between cells from the four timepoints. F) Scatterplot demonstrating that the correlation between the overall pseudotime trajectory and the vesicle-mediated transport trajectory is relatively tight across the range of values, and not driven by outliers. G) UMAP visualization of the expression of Rab15 and the progression of the vesicle-mediated transport trajectory show that Rab15 expression is relatively evenly found in the 48/72hr samples, but may not necessarily be found in every cell. H) Pseudotime trajectory of the interleukin signaling pathway shows a relatively simple path analogous to the main trajectory seen in Fig 3D. I) UMAP visualization of the expression of several key chemokine receptors and interleukin receptors confirms that most of these genes are downregulated upon stimulation with RANKL.

**Figure S9**-Switched genes along the osteoclast differentiation trajectory

A) Scatterplot mapping of all genes identified to display switch behavior (k magnitude greater than 1 and adjusted q-value < 0.05) along the overall pseudotime trajectory for osteoclast differentiation displayed in Fig 3B (ordered as t0). High intensity switched genes are defined as k >2, while low intensity switched genes have k <2. B) Histogram of the counts of switched genes along the pseudotime. Three primary peaks of switched genes can be identified, with some having switched almost immediately following stimulation, while others only shift at 5< t0 <10, corresponding to the shift between 24hr stimulated and 48/72hr stimulated cells. C) Enrichment dotplot showing the biological pathways (from Reactome) that the switched genes belong to. Notably, a number of elements of the vesicle-mediated transport and associated pathways displayed switch-like behavior. D) Focused analysis on the genes that switched in the window of 9< t0 <11, corresponding to the transition between 24hr-stimulated and 48/72hr stimulated cells, demonstrates that Rab15 switches within this timeframe. Interestingly, a number of mitochondrial proteins also shift with high intensity during this period, suggesting that metabolic changes may also occur at this timepoint.

**Figure S10-**Vesicular transport and osteoclast differentiation

A) Correlation network of the genes from the vesicular transport pathway generated using unstimulated and 24hr stimulated cells. Correlations with R > 0.6 are shown. B) Correlation network of the genes from the vesicular transport pathway generated using 48 and 72hr stimulated cells. Correlations with R > 0.6 are shown. C) Coexpression network of the genes most tightly co-expressed with Rab15. As expected, these include some of the genes found to be tightly associated with the overall pseudotime trajectory, such as Dap, Cry1, and Slc6a4. However, other factors such as the class II MHC invariant chain (Cd74) were also related and may suggest additional functional roles for Rab15. D) IGV visualization of the summed reads mapping to either Rab15 or Acp5 (encoding for Trap, a key enzyme for osteoclast function). Most of the reads for Rab15 map onto exon 7, the last exon of the gene. This expression pattern is a characteristic result of polyA-based approaches for mRNA capture in genes with lower absolute expression. (Genes with higher expression, such as Acp5, show additional reads that map to additional exons closer to the 5’ end and also have a higher likelihood of those sections being amplified by the NSRs.)

