## SupplementaryFIgures for "scSTATseq: Diminishing Technical Dropout Enables Core Transcriptome Recovery and Comprehensive Single-cell Trajectory Mapping"

**Figure S1-** scSTATseq workflow enhances transcriptome recovery

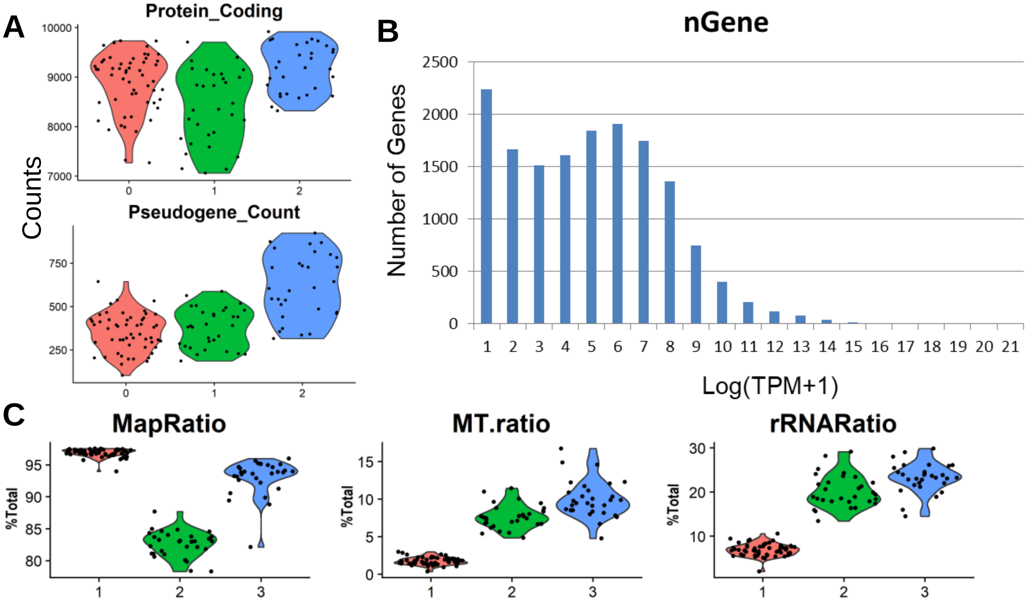

**Figure S2-** Cell cycle clustering of RAW cells

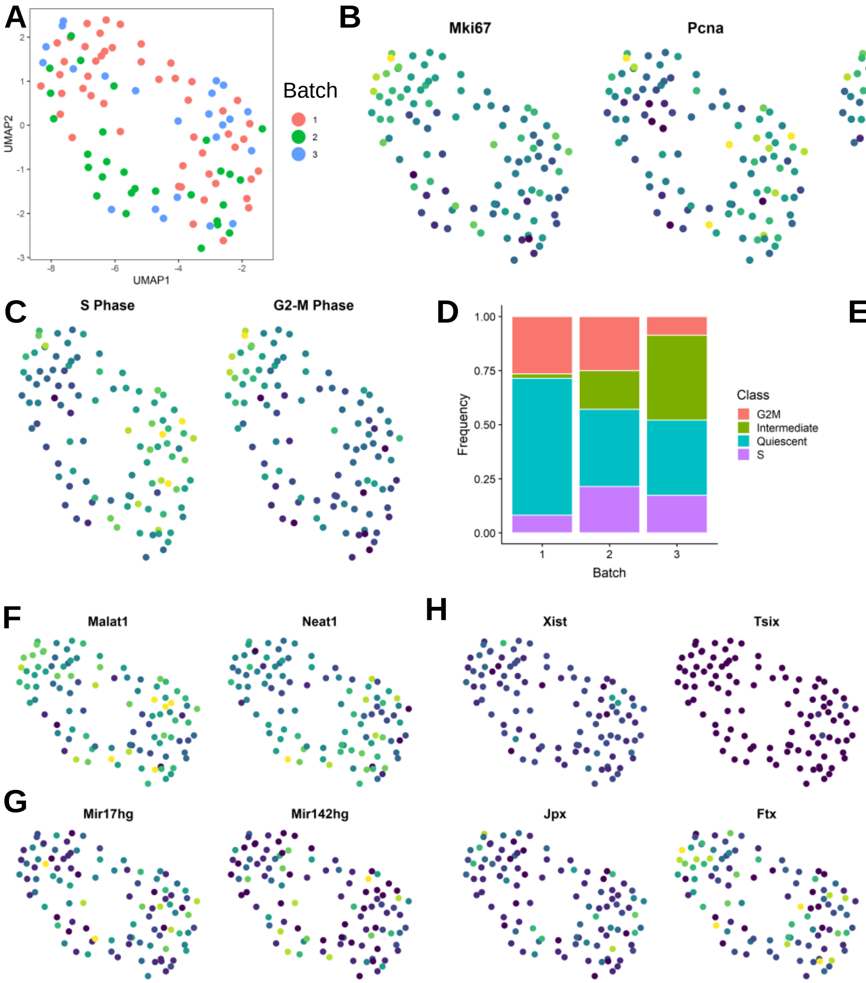

**Figure S3-** Broad correlation and distribution comparison

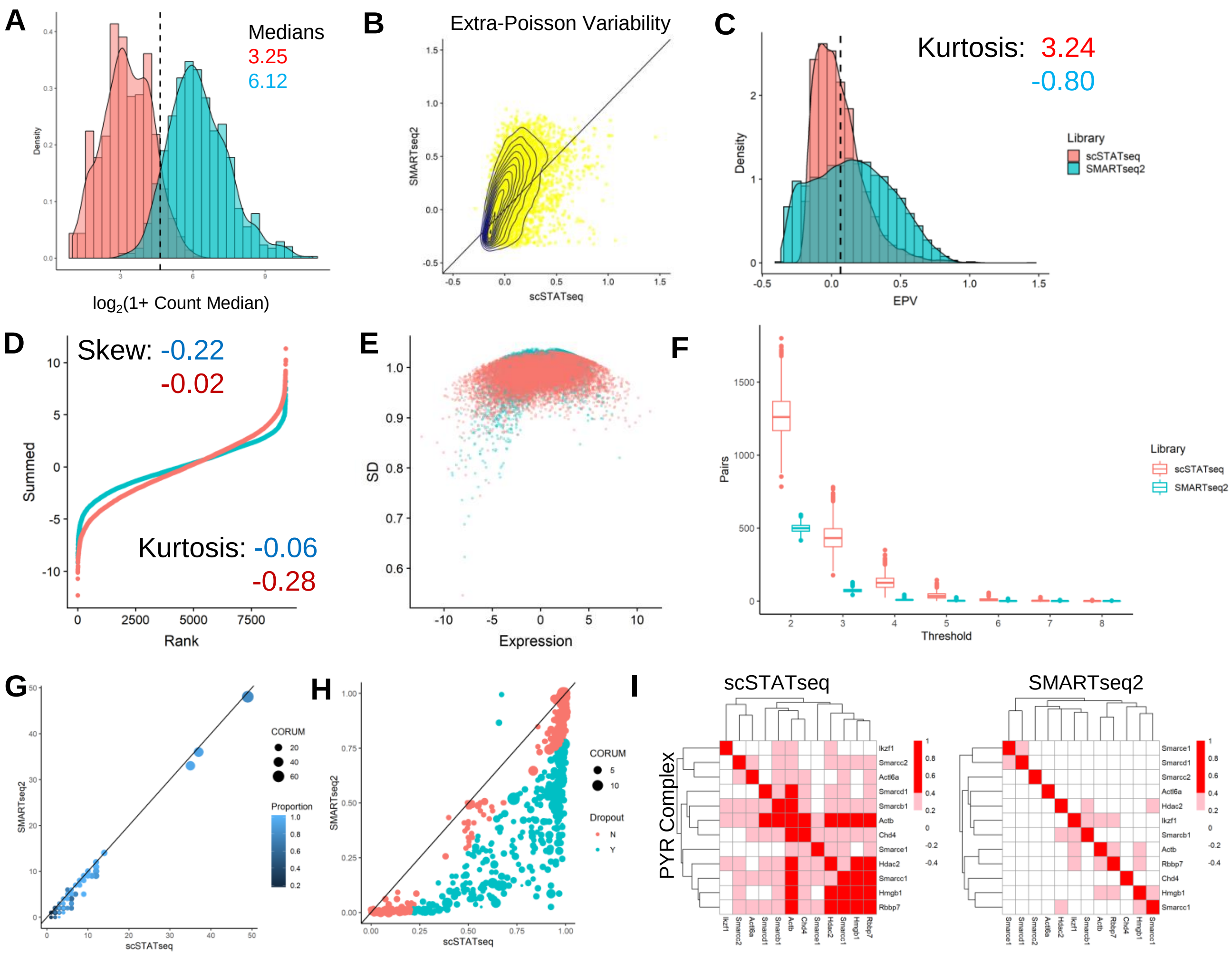

**Figure S4-** Correlated expression of KEGG pathways in scSTATseq data

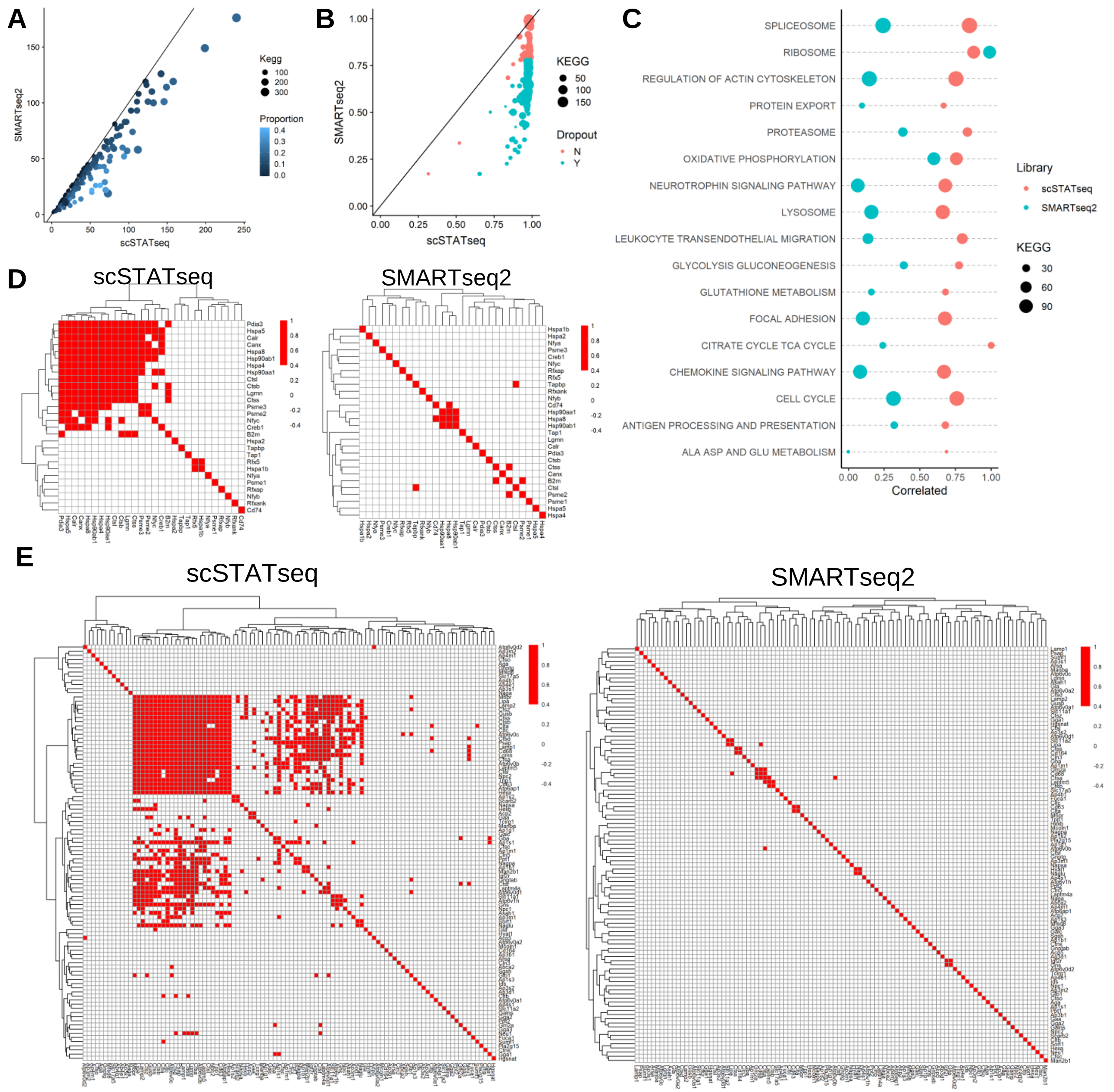

**Figure S5–** Core transcriptome of RAW cells

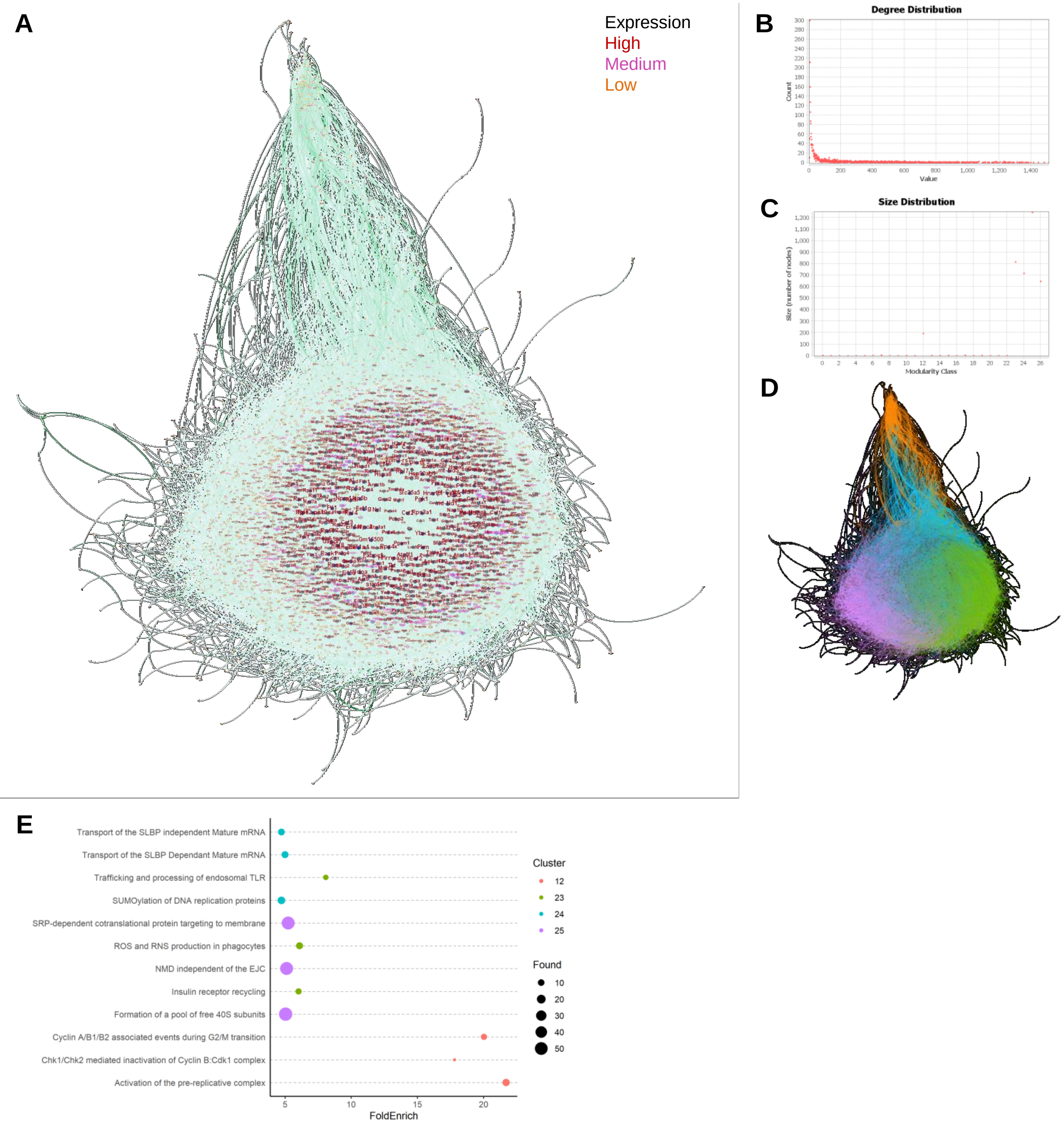

**Figure S6–** Immune-related core cluster

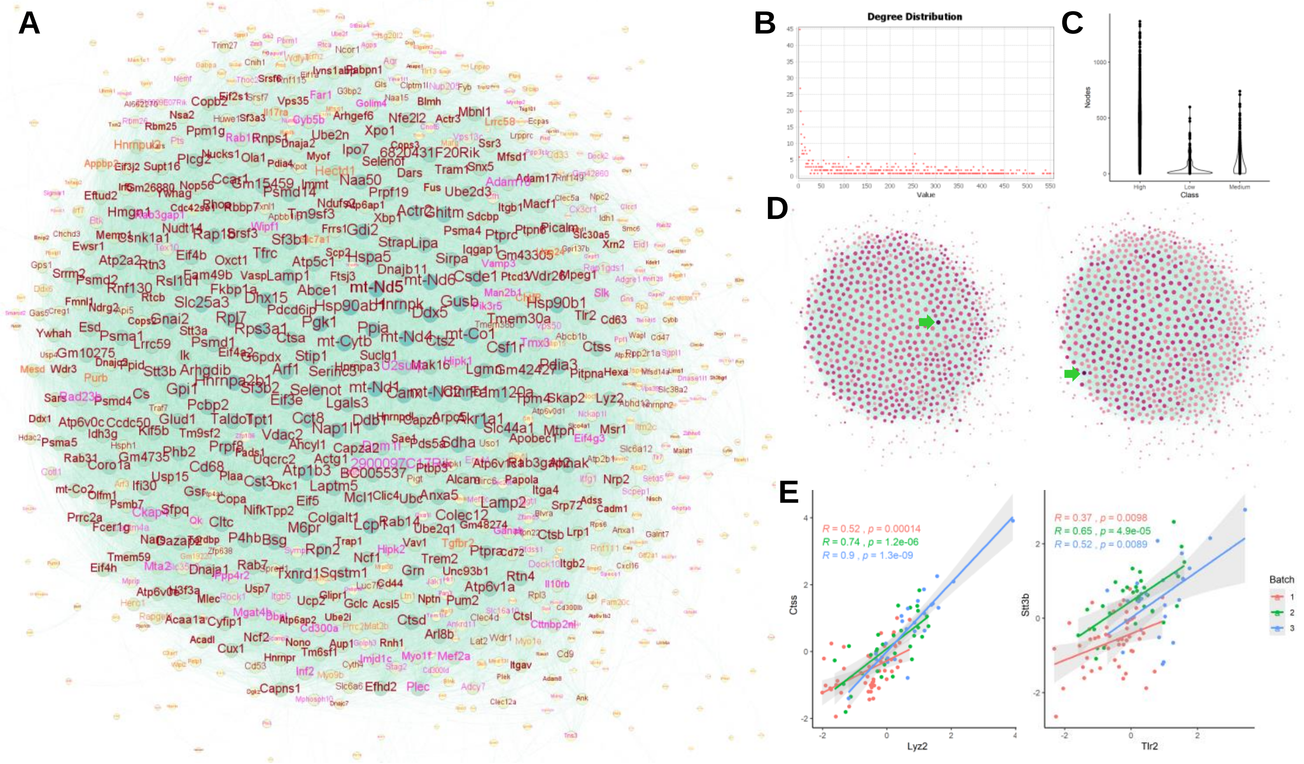

**Figure S7-** QC of osteoclast single cell sequencing

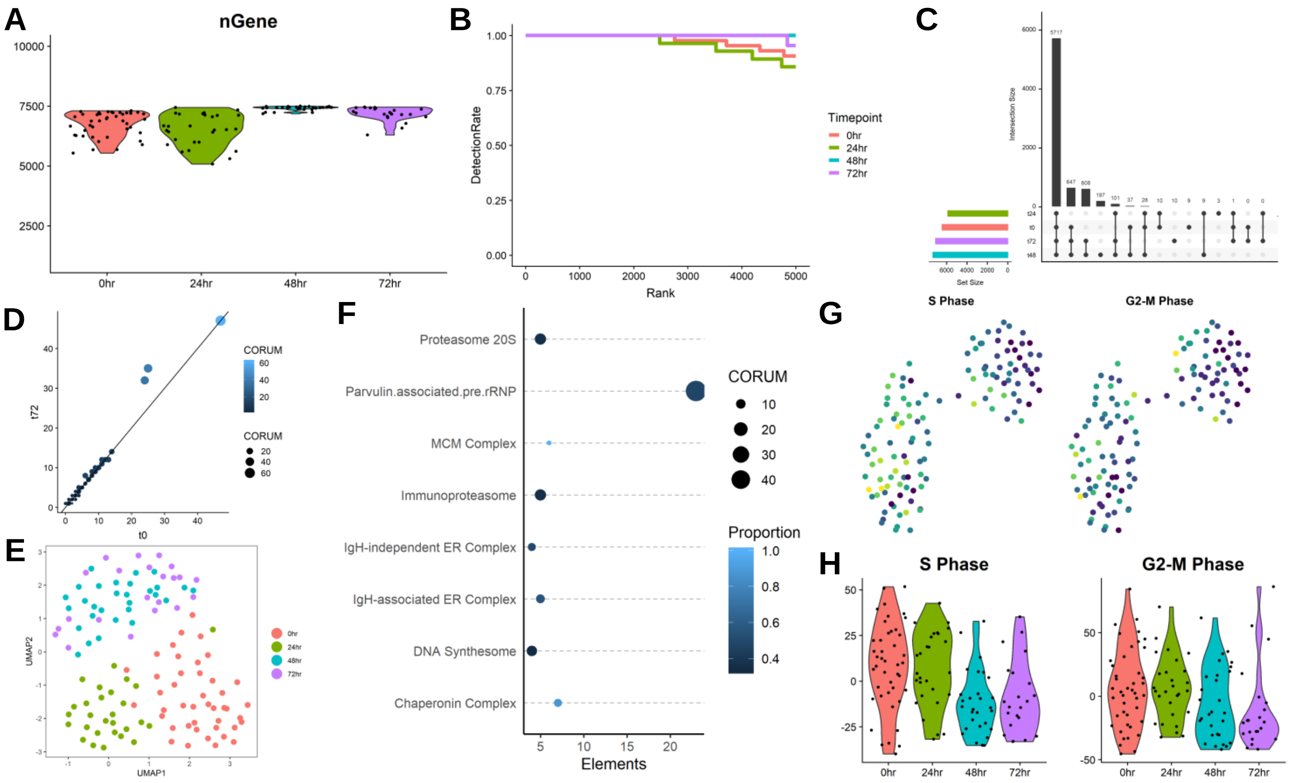

**Figure S8-** Branching trajectory of osteoclast differentiation

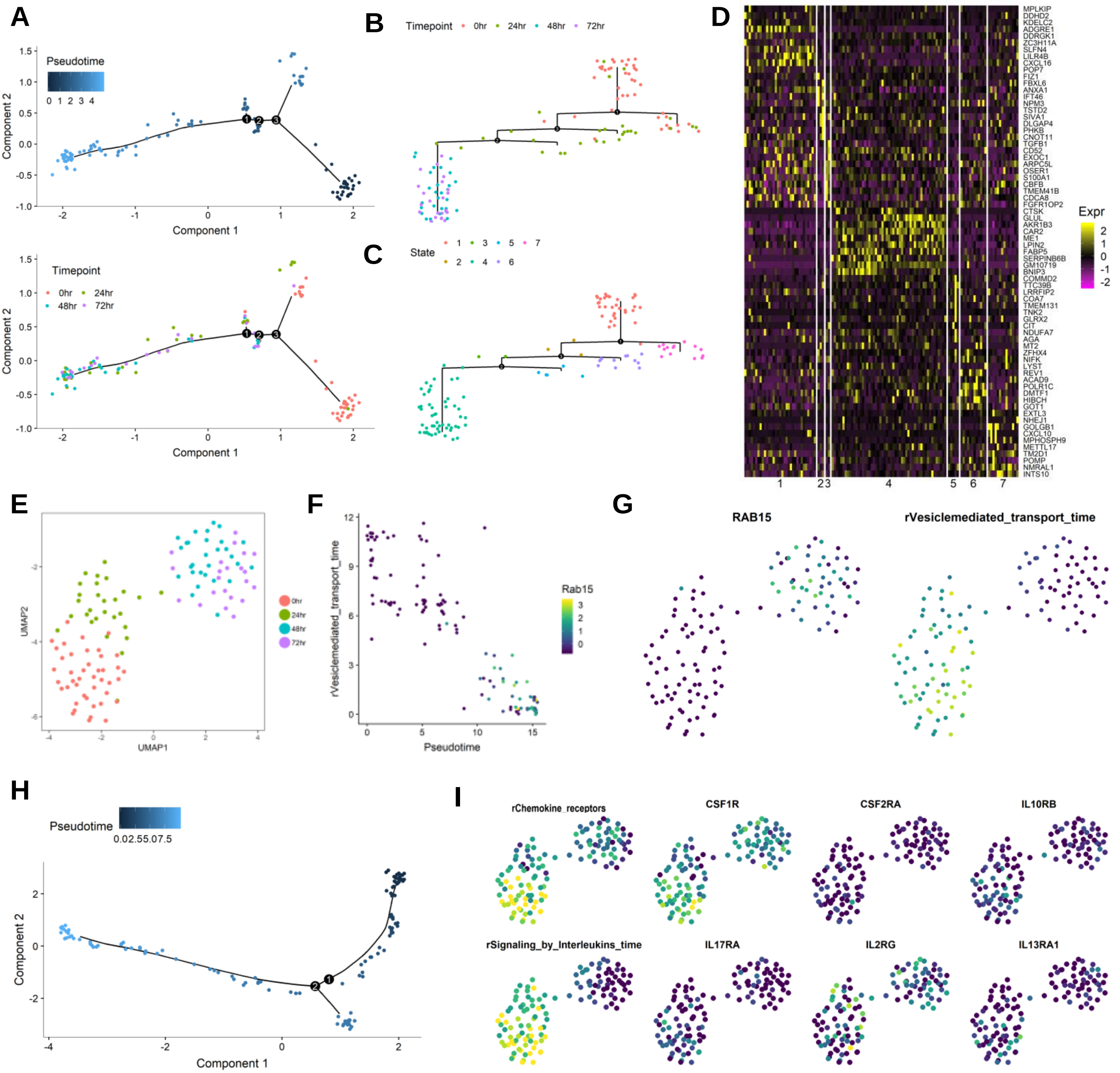

**Figure S9-** Switched genes along the osteoclast differentiation trajectory

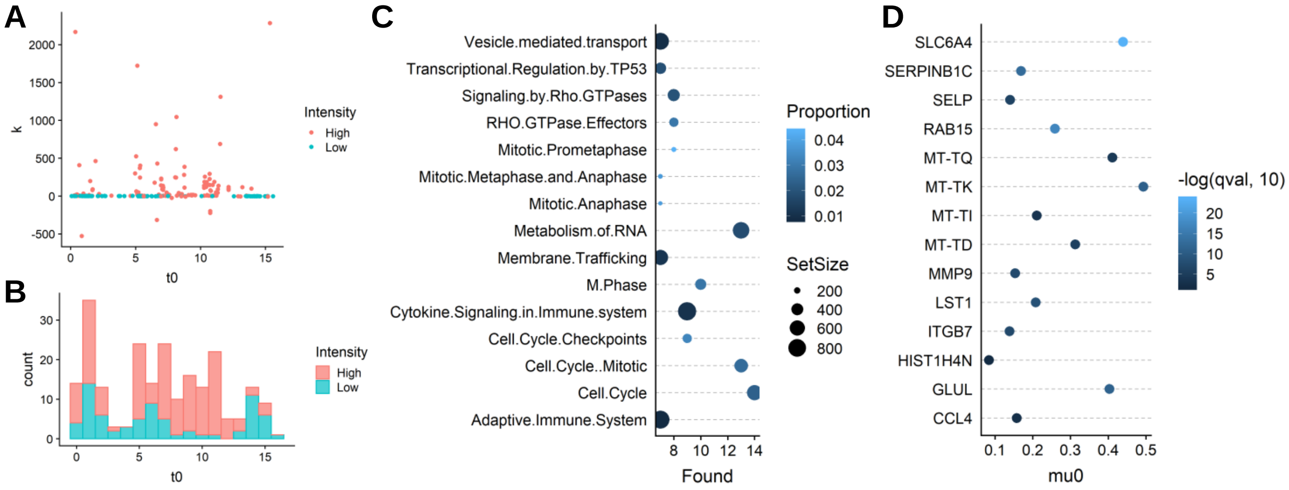

**Figure S10-** Vesicular transport and osteoclast differentiation

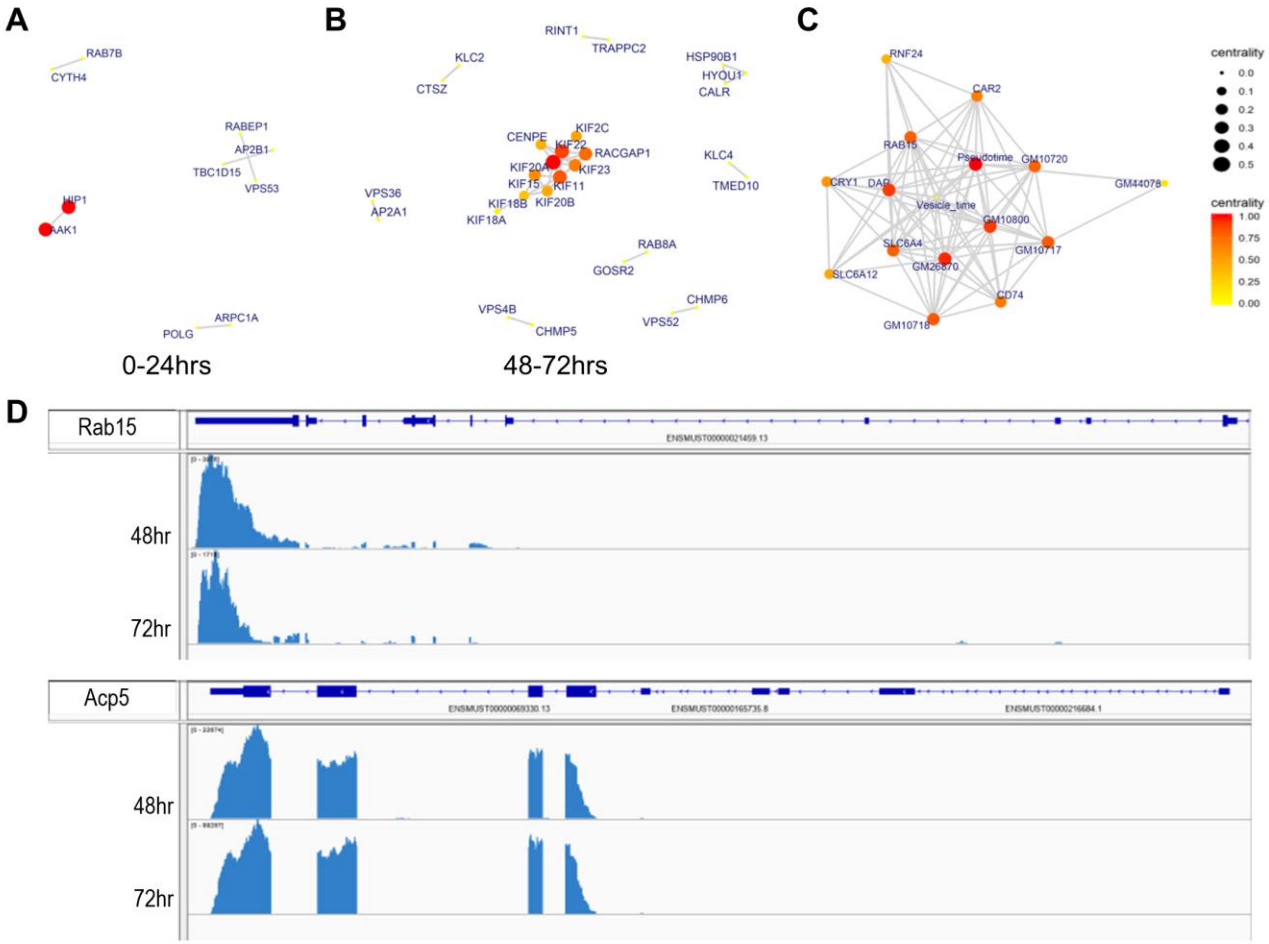
